# Development of LIBRA-seq for the Guinea Pig Model System as a Tool for the Evaluation of Antibody Responses to Multivalent HIV-1 Vaccines

**DOI:** 10.1101/2023.09.25.559382

**Authors:** Matthew J. Vukovich, Nagarajan Raju, Prudence Kgagudi, Nelia P. Manamela, Alexandra A. Abu-Shmais, Kathryn R. Gripenstraw, Perry T. Wasdin, Shaunna Shen, Bridget Dwyer, Jumana Akoad, Rebecca M. Lynch, David C. Montefiori, Simone I. Richardson, Penny L. Moore, Ivelin S. Georgiev

## Abstract

Consistent elicitation of serum antibody responses that neutralize diverse clades of HIV-1 remains a primary goal of HIV-1 vaccine research. Prior work has defined key features of soluble HIV-1 Envelope (Env) immunogen cocktails that influence the neutralization breadth and potency of multivalent vaccine-elicited antibody responses including the number of Env strains in the regimen. We designed immunization groups that consisted of different numbers of Env strains to be used in a cocktail immunization strategy: the smallest cocktail (group 2) consisted of a set of two Env strains, which were a subset of the three Env strains that made up group 3, which in turn were a subset of the six Env strains that made up group 4. Serum neutralizing titers were broadest in guinea pigs that were immunized with a cocktail of three Envs compared to cocktails of two and six, suggesting that multivalent Env immunization provides a benefit but may be detrimental when the cocktail size is too large. We then adapted the LIBRA-seq platform for antibody discovery to be compatible with guinea pigs, and isolated several tier 2 neutralizing monoclonal antibodies. Three antibodies isolated from two separate guinea pigs were similar in their gene usage and CDR3s, establishing evidence for a guinea pig public clonotype elicited through vaccination. Taken together, this work investigated multivalent HIV-1 Env immunization strategies and provides a novel methodology for screening guinea pig B cell receptor antigen specificity at a high throughput level using LIBRA-seq.

**IMPORTANCE:** Multivalent vaccination with soluble Env immunogens is at the forefront of HIV-1 vaccination strategies, but little is known about the influence of the number of Env strains included in vaccine cocktails. Our results suggest that adding more strains is sometimes beneficial but may be detrimental when the number of strains is too high. Additionally, we adapted the LIBRA-seq platform to be compatible with guinea pig samples and isolated several tier 2 neutralizing monoclonal antibodies, some of which share V and J gene usage and >80% CDR3 identity, thus establishing the existence of public clonotypes in guinea pigs elicited through vaccination.

## INTRODUCTION

Although significant progress has been made in the field of HIV-1 vaccine design, an effective vaccine capable of eliciting broadly neutralizing antibody responses is yet to be developed. The HIV-1 Envelope protein (Env) is the sole target of neutralizing antibodies and employs several strategies to evade the immune response, including but not limited to extensive glycosylation to shield vulnerable protein epitopes, presentation of immunodominant non-neutralizing or strain-specific neutralizing epitopes, conformational masking, and vast amino acid sequence diversity [1–7]. Despite these challenges, some chronically infected individuals develop broadly neutralizing serum primarily attributed to the development of broadly neutralizing antibodies (bNAbs) [8–14]. Efforts to recapitulate a bNAb response through vaccination have proven extremely difficult, largely due to the unique features of bNAbs including long CDRH3s, improbable mutation sets, and rare bNAb-producing B cell lineages available for activation [1, 15].

Guinea pigs have proven to be an excellent model system for studying vaccine efficacy, especially in the field of HIV-1 vaccine design[16–19]. Although there is a long history of using guinea pigs in vaccine studies, there is an inadequate toolkit available to study the elicited antibody response at the monoclonal level in a high-throughput and reproducible manner. Most vaccine studies with guinea pigs rely on measuring the polyclonal neutralizing response as a readout of vaccine efficacy, which can miss the nuanced contributions of rare desirable antibodies [17–22]. Currently available antibody sequencing techniques for guinea pigs rely on single-cell sorting into 96-well plates, with little information that can help down-select candidate antibodies for expression and purification [23]. To this end, we sought to adapt our LIBRA-seq (linking B cell receptor to antigen specificity through sequencing) platform to be compatible with guinea pigs, thus allowing for a high-throughput technology to assess the antigen specificity of guinea pig B cells against potentially large numbers of antigen variants, while also providing a prioritization metric (the LIBRA-seq Score) to down-select candidate antibodies for validation. This technology will be especially beneficial for deconvoluting the antigen specificity patterns in antibody responses to multivalent vaccines (cocktails of immunogens).

Researchers have made significant progress through multivalent vaccination approaches targeted against HIV-1. Inclusion of more Env strains in a vaccine has been shown to increase the elicited neutralization breadth [17, 18, 22, 24]. More recent work systematically defined Env amino acid signatures associated with sensitivity to bNAbs and appended these amino acid motifs onto Env immunogens, termed signature-based epitope targeting (SET) immunogens [18]. When combined with an immunogen engineered to display higher epitope diversity, SET immunogens achieved remarkable breadth in a guinea pig model. To further investigate important factors of multivalent Env immunization regimens, we explored the effect of the number of Env strains included in cocktail immunizations. We computationally selected different Env cocktail sizes while simultaneously optimizing bNAb neutralization sensitivity and Env amino acid sequence diversity. Env immunogens that are missing conserved glycans tend to elicit strain-specific neutralizing antibodies that target the glycan holes but lack the ability to neutralize virus isolates that have an intact glycan shield [2, 25]. To mitigate bias introduced by glycan holes, Env strains were selected based on the presence of potential N-linked glycosylation sites (PNGS). We immunized guinea pigs with computationally selected cocktails of two, three, or six different Env strains and evaluated the serum neutralizing antibody response at the polyclonal level. As a proof of concept, we adapted and leveraged our LIBRA-seq platform to interrogate the B cell repertoire of the two guinea pigs with the greatest polyclonal breadth of neutralization (gp109 and gp112), and isolated several hundred B cells with B cell receptor (BCR) sequences that were predicted to bind to diverse strains of Env based on their LIBRA-seq scores. We selected thirty antibodies for expression and validated binding for twenty-seven of these antibodies, of which six neutralized tier 2 pseudoviruses. Notably, three of these antibodies (two from gp112 and one from gp109) used the same germline V and J-genes and remarkably similar CDR3 regions for both the heavy and light chains, establishing the existence of vaccine-elicited guinea pig public antibody clonotypes.

## RESULTS

### Immunization groups

The vaccine trial was composed of four groups. Group 1 consisted of a pair of strains (BG505 T332N and CZA97.012) that in prior studies have not elicited a robust neutralizing antibody response in cocktail immunizations, serving as a baseline control. Groups 2-4 consisted of strains that were selected using an algorithm that optimized for glycan shield coverage, sensitivity to known bNAb classes, and Env sequence diversity (as described in methods), but for different numbers of strains in each cocktail. Group 2 consisted of two strains (5768.04 and 286.36), Group 3 consisted of three strains (5768.04, 286.36, and KNH1209.18), and Group 4 consisted of six strains (5768.04, 286.36, KNH1209.18, HT593.1, MB539.2B7, and DU172.17) (Figure 1A).

**Figure 1:**
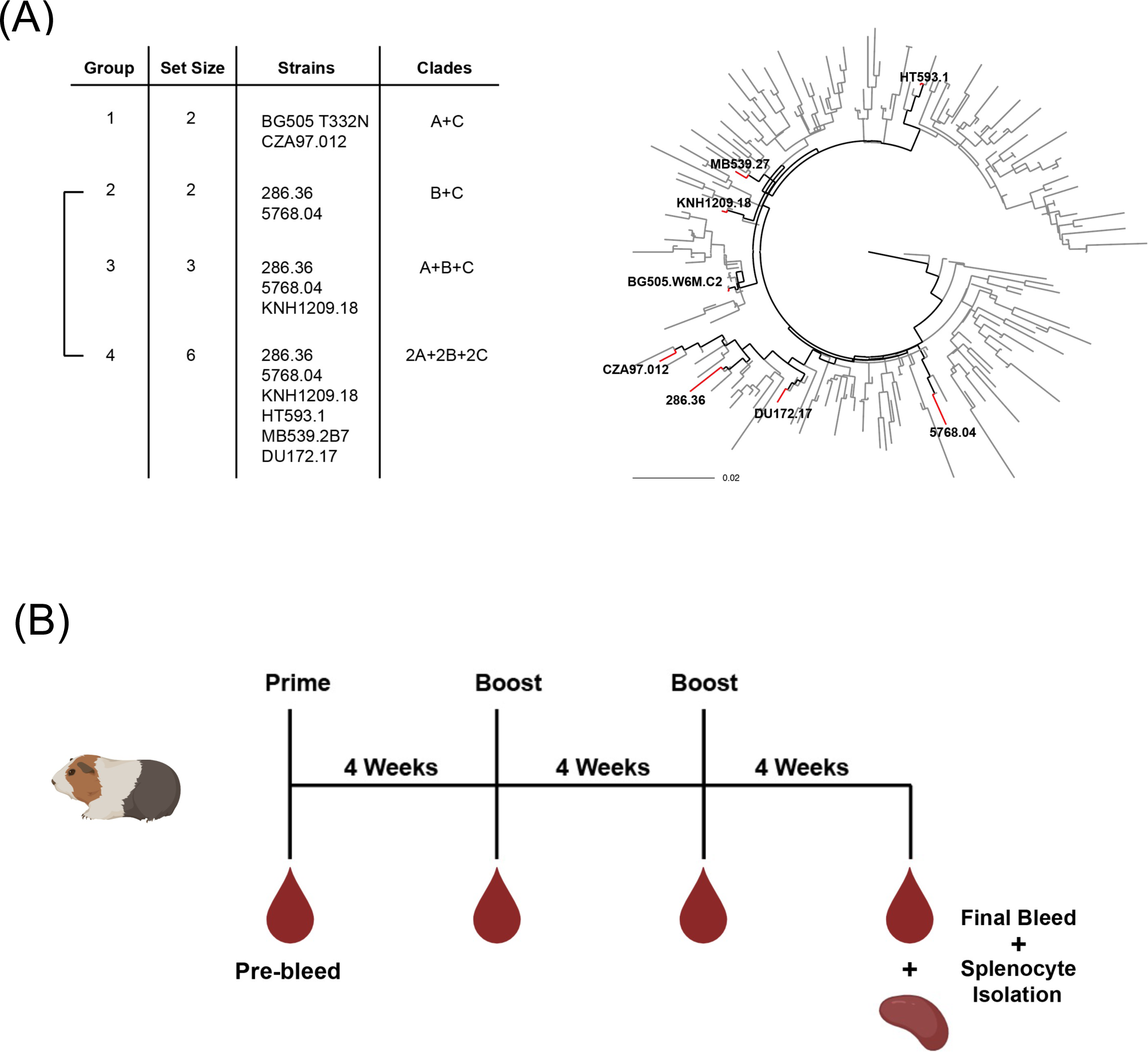
Vaccine groups were designed to evaluate the effect of including different numbers of Env strains in a cocktail immunization strategy. 1A: Vaccine groups were composed of two strains (group 1 and 2), three strains (group 3), or four strains (group 4). Group 1 consisted of two strains that have traditionally been used in the past. Envs in groups 2-4 were chosen to primarily assess the effect of having different numbers of Env strains included in the vaccine and were chosen by maximizing Env amino acid sequence diversity, as well as selecting for strains with intact glycan shields and high sensitivity to all major bNAb classes. 1B: Guinea pigs were immunized with 100µg total SOSIP.DS.644 immunogen at three intervals four weeks apart. Final bleeds and splenocytes were collected four weeks after the final dose.

### Immunogenicity of the multivalent vaccine cocktails

Guinea pigs were vaccinated intramuscularly in the quadriceps with 100 μg total Env immunogen formulated with CpG/Emulsigen at weeks 0, 4, and 8 (Figure 1B). For each injection, half of the formulation was injected into the left quadricep, and the other half was injected into the right quadricep. Serum was collected at each time point prior to injections. Guinea pigs were exsanguinated 4 weeks after the final injection, and splenocytes were isolated for downstream antigen-specific B cell sorting. All groups showed a robust antibody response towards autologous and heterologous strains of Env trimer by ELISA (Figure S1). In addition to serum antibody binding, we evaluated the ability of antibodies in the serum to mediate antibody-dependent cellular phagocytosis (ADCP) in a bead-based assay and found that serum from all groups exerted ADCP to autologous and heterologous antigens to varying degrees (Figure S2). Next, we evaluated the neutralizing antibody response using serum from the final timepoints against a large panel of HIV-1 pseudoviruses that included eight vaccine strains and eleven global panel strains (Figure 2). Overall, group 3 exhibited the broadest levels of serum neutralization with a median number of detectable responses against 5 pseudoviruses (26% breadth) compared to a range of 3-3.5 (16-18% breadth) for the other groups (Figure 2B). Notably, 5/6 animals in group 3 had detectable neutralizing responses against at least 26% of the full virus panel, and 2/6 animals neutralized at least 58% of the viruses. In comparison, only 1/5 animals in group 2 and 2/5 animals in group 4 showed neutralizing responses against at least 26% of the viruses. Interestingly, when taking into account the maximum and minimum number of detectable responses, group 4 (with a cocktail size of six) underperformed compared to all other groups with smaller cocktail sizes. These data suggest that increasing the number of diverse strains in a cocktail regimen has the potential to expand neutralization breadth, but that adding more strains is not always beneficial.

**Figure 2:**
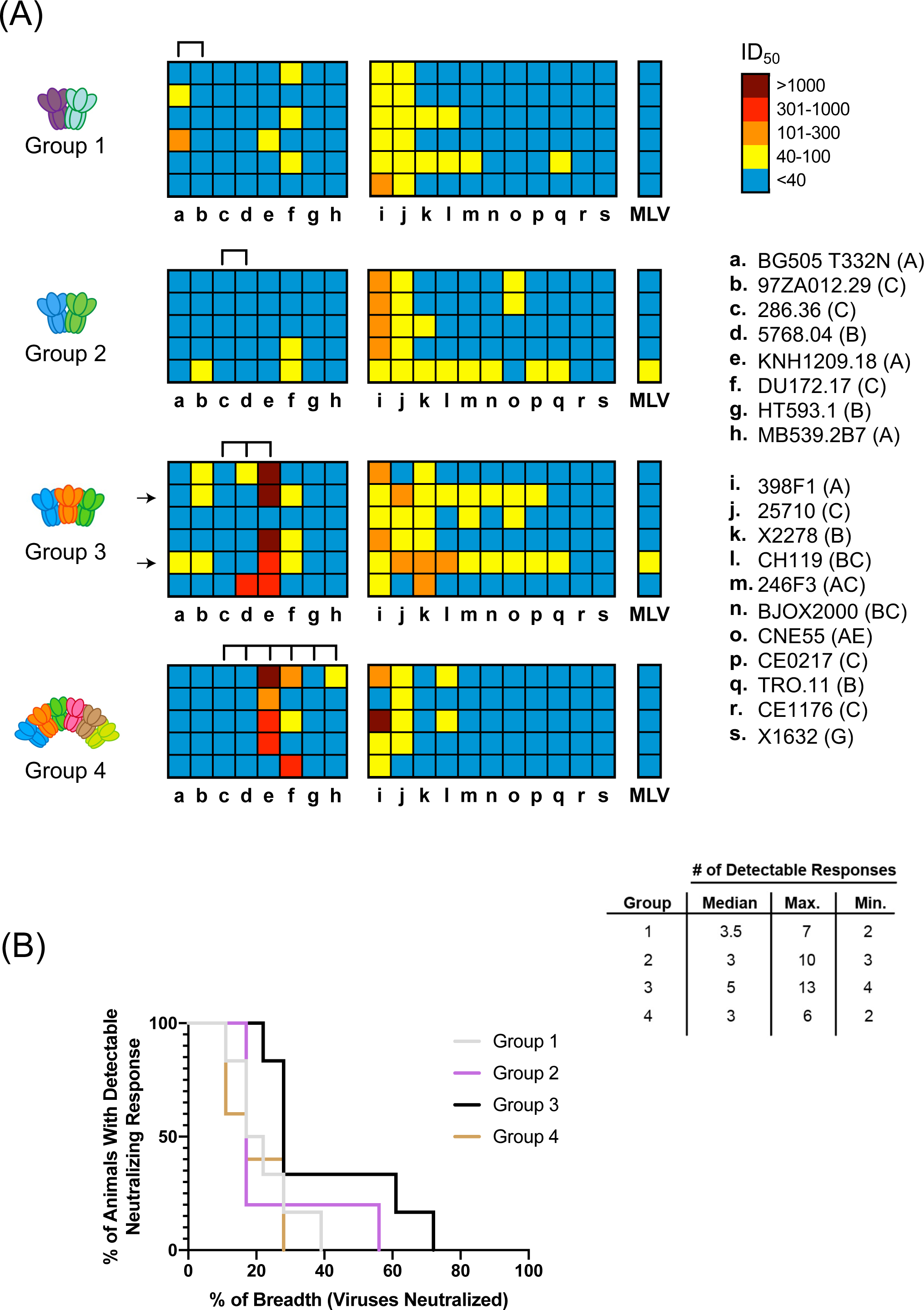
Serum neutralization data for vaccine groups. 2A: Guinea pig sera was tested for neutralization against HIV-1 pseudoviruses displaying Envs from vaccine strains (a-h) and the standard global panel (i-s) in the TZM-bl cell neutralization assay. Each row displays serum neutralization data from one guinea pig and each column represents neutralization data against a given strain of pseudovirus. ID_50_ values are reported as colors ranging from no neutralization detected (blue) to high potency neutralization (dark red). Arrows denote guinea pigs 109 (top) and 112 (bottom), from which splenocytes were used for monoclonal antibody isolation. 2B: Comparison of breadth between groups as a function of the proportion of guinea pigs in each group that exhibit greater than or equal to that breadth denoted on the x-axis.

### Isolation and characterization of tier 2 neutralizing monoclonal guinea pig antibodies with LIBRA-seq

We next sought to adapt LIBRA-seq for use with guinea pig splenocytes to isolate tier 2 neutralizing antigen-specific monoclonal antibodies from gp109 and gp112 (indicated with arrows in Figure 2A). LIBRA-seq is a single-cell sequencing technology for high-throughput screening of B cells against a diverse antigen panel labeled with oligonucleotide barcodes [26]. We first designed a flow panel that allowed us to isolate antigen-positive IgG-expressing B cells. We isolated IgG^Positive^ IgM^Negative^ cells to bias towards IgG class-switched B cells, followed by isolation of antigen-positive B cells (Figure 3A). Our LIBRA-seq antigen panel consisted of 3 autologous Envs (286.36, 5768.04, and KNH1209.18) that were part of the group 3 immunization cocktail, 2 heterologous Envs (HT593.1 and BG505 T332N), a gp120-based CD4 binding site-specific antigen (RSC3), and a negative control antigen (CA/09 H1), all of which were biotinylated and conjugated with distinct oligonucleotide barcodes to allow for, respectively, staining with PE-streptavidin and antigen identification through sequencing (Figure 3B). A critical step in single-cell sequencing to obtain paired heavy and light BCR chains is target enrichment following cDNA generation. The inner and outer reverse primers were generated based on previous work with antigen-specific B cell sorting with guinea pig samples [23], whereas the inner and outer forward primers are not species-specific, and thus did not require alteration (Figure 3B).

**Figure 3:**
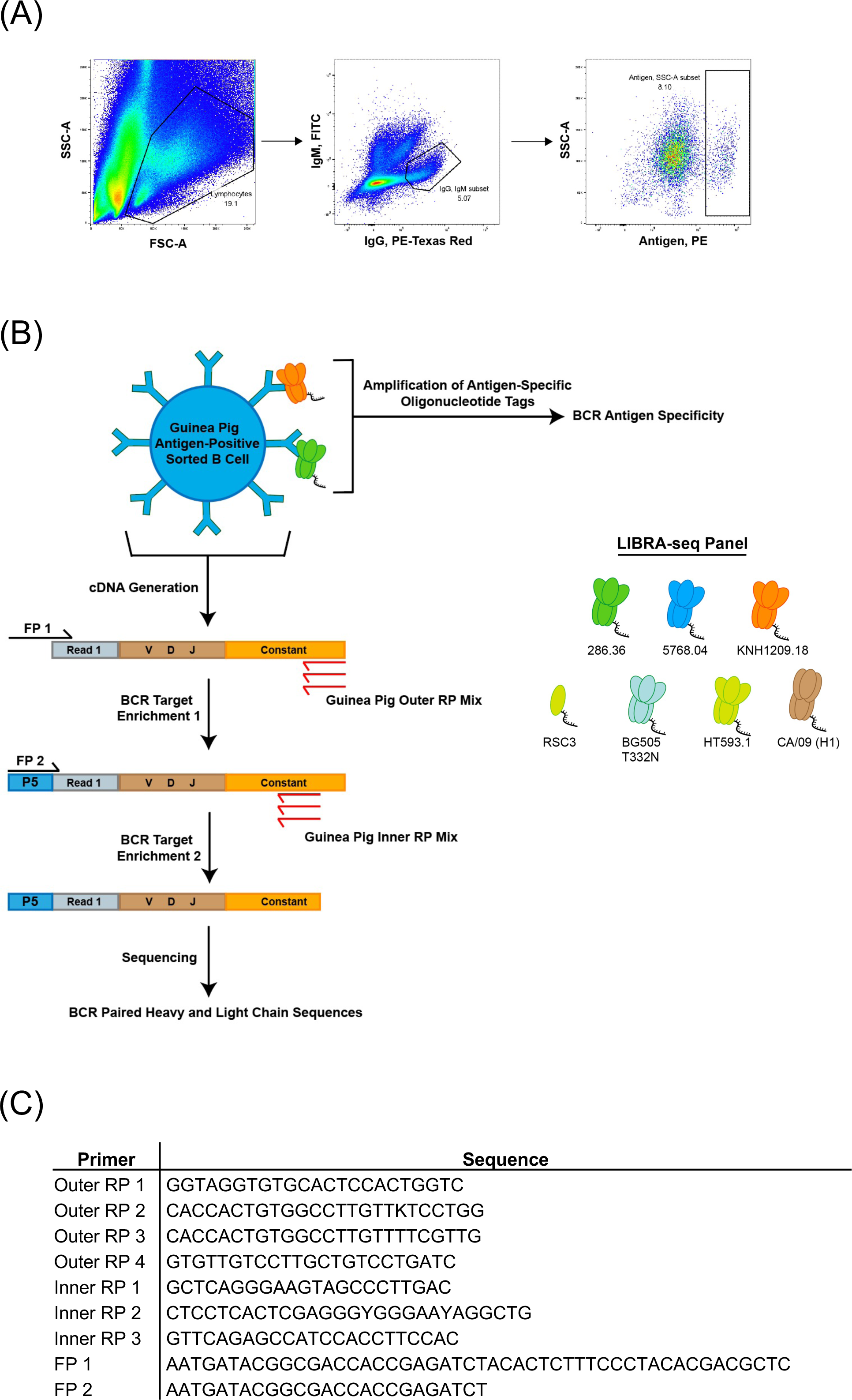
Development of LIBRA-seq with guinea pig B cells. 3A: flow panel was designed to isolate antigen-positive guinea pig B cells from splenocytes obtained from guinea pig 109 and 112, chosen based on their broad serum neutralization profiles. Lymphocytes were gated out based on size, followed by live cells (data not shown). Cells with high IgG expression and negative for IgM expression were then gated followed by antigen positive B cells. Data shown is from guinea pig 112. 3B: The LIBRA-seq workflow was adapted for use in guinea pigs by modifying BCR sequencing steps. Primers shown in red indicate guinea pig-specific reverse primers that anneal to the constant regions of the guinea pig IgG genes. These custom reverse primers were mixed with existing forward primers specific to sequences appended during single cell sequencing using the Chromium Next GEM Single Cell V(D)J Reagent Kit. Following target enrichment single cell sequencing was carried out as usual. The LIBRA-seq antigen screening panel is shown to the right and consisted of 5 Envs, the RSC3 antigen, and an influenza HA CA/09 (negative control). 3C: Sequences of the primers used in guinea pig LIBRA-seq.

From the LIBRA-seq output we obtained a total of 277 cells from gp109, and 108 cells from gp112. From these we selected thirty paired heavy and light chain sequences to express as humanized IgG1 antibodies (five from gp109 and twenty-five from gp112) based on antigen specificity scores for Envs in the LIBRA-seq panel (Figure 4A). V-gene usage as well as CDR3 sequence and length for all thirty antibodies are reported (Figure S3A). Of the thirty antibodies, twenty-seven bound to at least one Env antigen, and several bound to multiple diverse strains. In total, 90% of the LIBRA-seq-identified cells were confirmed to be HIV-specific. Next, we evaluated the antibodies for neutralization against a panel of autologous strains of tier 2 HIV-1 pseudoviruses (286.36, 5768.04, and KNH1209.18) in the TZM-bl cell neutralization assay. A total of six antibodies were identified as neutralizing, with three potently neutralizing the KNH1209.18 strain and three neutralizing the 5768.04 strain with low potency (Figure 4B). The six neutralizing antibodies were found to be strain-specific as they did not neutralize any other strains in the vaccine panel nor a global panel, with the exception of antibody 112-9 which showed weak neutralization against CNE55 (Figure S3B). Sequence comparison of the three antibodies that neutralized the KNH1209.18 strain (two from gp112 and one from gp109) revealed the use of the same germline genes and high CDR3 identity for both the heavy and light chains, suggesting that we were able to recover a vaccine-elicited public guinea pig clonotype (Figure 4C).

**Figure 4:**
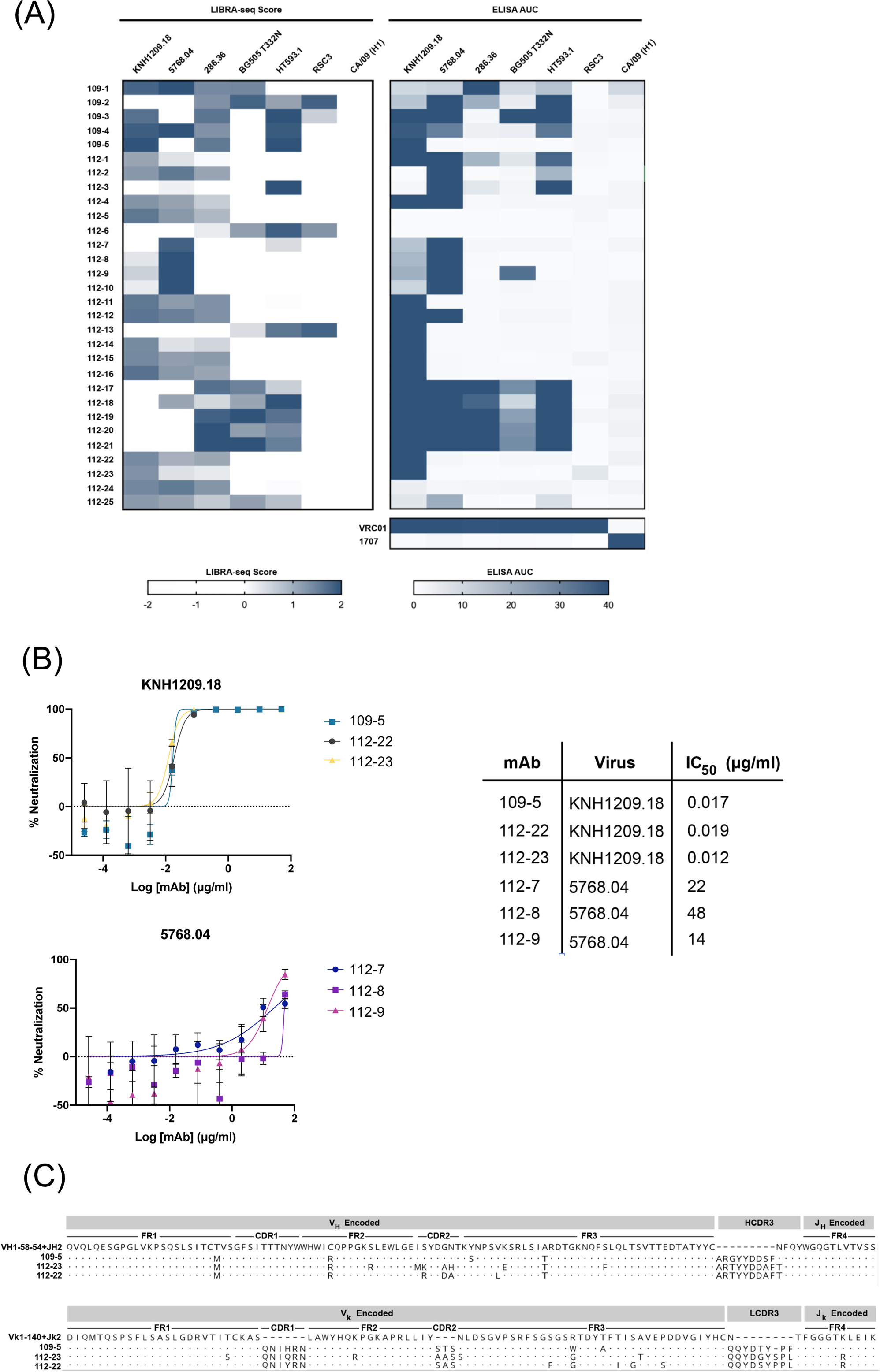
Identification and validation of tier 2 neutralizing monoclonal antibodies from vaccinated guinea pigs using LIBRA-seq. 4A: The LIBRA-seq output showing prioritized antibodies from guinea pig samples with predicted antigen specificities is shown. Each row consists of one B cell with the LIBRA-seq scores for each antigen, ranging from low (white) to high (blue), as well as ELISA area under the curve (AUC) with the same color scheme. Antibodies isolated from guinea pig 109 have a 109 before their number, and antibodies from guinea pig 112 have a 112 before their number. 4B: Six neutralizing antibodies were identified from the positive hits. The virus strain is indicated by the title for each curve. IC_50_ values are displayed on the chart to the right of the curves. 4C: Sequence analysis revealed a guinea pig public clonotype. Antibodies 109-5 (from guinea pig 109) along with antibodies 112-22 and 112-23 (from guinea pig 112) share the same variable and joining gene usage and have >70% CDR3 identity.

## DISCUSSION

We investigated the effect of the number of Env strains included in a multivalent HIV-1 vaccine on the elicited serum neutralization breadth in guinea pigs. In parallel, we adapted our LIBRA-seq platform to be compatible with guinea pig vaccine samples, and applied this technology to the vaccine study described here, as a proof of concept, thus expanding the toolkit for studying vaccine-elicited antibody responses at the monoclonal level. The Env strains used in the immunization cocktails were selected using criteria that have been shown to be critical in eliciting broadly neutralizing antibody responses, including glycan shield coverage, bNAb neutralization sensitivity, and Env amino acid sequence diversity. Immunization group 3, immunized with a cocktail of three Envs, exhibited the broadest neutralizing serum of all the groups. This could suggest that in the context of multivalent HIV-1 vaccines, larger cocktail sizes are not always better, as was evidenced with the results for group 4, immunized with a cocktail of six Envs that included all Envs from group 3 along with three additional diverse Envs.

It is interesting, however, to further dissect the differences in neutralizing serum responses for the three immunization groups, given that the Env strains in group 2 are a subset of the Env strains in group 3, which in turn are a subset of the Env strains in group 4. None of the animals from group 2 showed neutralization against the two autologous viruses (286.36 and 5768.04). In contrast, two animals from group 3 (that also included KNH1209.18 in addition to the other two viruses from group 2) were able to neutralize 5768.04, with one of the animals reaching high potency (ID_50_ of >300). These observations suggest that the addition of an additional strain can modulate the immune response toward neutralization against an autologous strain for which neutralization was otherwise not observed. Compared to group 4, a larger number of animals from the group 3 and group 2 sets were capable of neutralizing higher fractions of the tested viruses. While this is a single example with limited data, it is also interesting to note that adding more strains to multivalent vaccines does not necessarily result in improved neutralization breadth, compared to groups with smaller sizes (group 3 was better than both the group 4 and group 2). Together, these results indicate that the selection of strains for incorporation into multivalent vaccines can indeed impact neutralization breadth. We note that the results with the group 3 set are the most promising among the four multivalent vaccines tested, although there are certainly areas that can be improved upon, including the elicitation of autologous neutralization against all strains that are part of the multivalent vaccine, consistent neutralization breadth among all animals, and increased overall potency against heterologous viruses. Further investigation of how to systematically optimize multivalent HIV-1 vaccines, including both strain selection and optimization, therefore will be valuable.

The ability to characterize the vaccine-elicited antibody response at the monoclonal level is paramount to understanding vaccine trials and designing next-generation vaccines. We adapted our LIBRA-seq technology to be compatible with guinea pig samples, which allowed us to probe the antigen specificity of the guinea pig B cell repertoire at unprecedented levels. We validated LIBRA-seq on guinea pig samples using splenocytes from the two guinea pigs in group 3 that had the broadest polyclonal neutralization, gp109 and gp112. We used a diverse panel of Env antigens in the LIBRA-seq screening library to help prioritize antibody leads for further characterization. Several neutralizing antibodies were identified among those tested for neutralization. Of note, our serum neutralization assays failed to detect any neutralization against the 5768.04 strain, for which three of the monoclonal antibodies showed modest neutralization ability. This emphasizes the importance of being able to isolate and characterize antibodies at the monoclonal level, as it allows us to identify rare antibodies that may not be reflected in the signals observed with polyclonal assays. Although previous HIV-1 vaccine studies with guinea pigs have reported elicitation of polyclonal sera capable of neutralizing tier 2 pseudoviruses, most did not isolate monoclonal antibodies [18, 20]. In this study, we utilized reverse primers specific for the IgG guinea pig BCR genes for target enrichment during sequencing. Additional isotypes may play a key role in the adaptive immune response to HIV-1 and other pathogens; as such, it will be of interest in future studies to design additional primers to allow for sequencing of the entire guinea pig BCR repertoire. On a similar note, the currently available guinea pig reference genome is not yet suitable for computational pipelines such as Cell Ranger. In this study we utilized a human BCR reference, which likely only allowed the recovery of a subset of guinea pig BCR sequences (those that map close enough to human BCR sequences). It will be beneficial to develop a more complete guinea pig reference to allow for full interrogation of the guinea pig repertoire.

Sequence analysis of the three antibodies that potently neutralize the KNH1209.18 strain (109-5, 112-22, and 112-23) revealed the existence of vaccine-elicited public clonotypes in guinea pigs. In addition to the similar point mutations present on the heavy chains, a large insertion of 6 similar amino acids on the light chain of all three antibodies was identified (Figure 4C). Since it is unlikely that both guinea pigs developed these same features through mutations, especially the six amino acid insertion in the light chain, we suggest that there may be germline BCR genes in the guinea pig repertoire that have yet to be described. While there are similarities between the three antibodies, there were multiple residue positions with divergent mutations in the different antibodies, likely indicating that antigen-driven evolution of the antibodies occurred.

We note that guinea pigs are a popular model for assessing vaccine responses, and that multivalent vaccine cocktails are of interest for a variety of pathogens of biomedical significance, including influenza, hepatitis C, SARS-CoV-2, and HIV-1 among others. As such, the LIBRA-seq technology can play an important role as a general platform for vaccine development and evaluation.

## ACKNOWLEDGMENTS

We thank all members of the Georgiev lab for their support and feedback. We thank David Flaherty, Olivia Murfield, Emma McLaughlin, and Brittany Matlock from the VUMC Flow Cytometry Shared Resource, for their help with cell sorting. The VUMC Flow Cytometry Shared Resource is supported by the Vanderbilt Ingram Cancer Center (P30 CA68485) and the Vanderbilt Digestive Disease Research Center (DK058404). We thank Angela Jones, Jamie Roberson, Latha Raju, and Lana Olson with the Vanderbilt Technologies for Advanced Genomics Core (VANTAGE) for providing technical assistance with library production and sequencing. VANTAGE is supported in part by CTSA (5UL1 RR024975-03), the Vanderbilt-Ingram Cancer Center (P30 CA68485), the Vanderbilt vision Center (P30 EY08126), and NIH/NCRR (G20 RR030956). We thank Christopher Aiken for his help with preparing pseudoviruses. We thank Tandile Hermanus for technical assistance with functional characterization of antibodies. For work described in this manuscript, I.S.G., M.J.V, N.R, A.A.A., K.R.G, and P.T.W. were supported in part by NIH R01AI175245 (to I.S.G.). A.A.A. was supported in part by NIH grant T32 (5T32AI112541-07); P.T.W was supported in part by NIEHS grant T15 (T15LM007450-19). P.L.M. is supported by the South African Research Chairs Initiative of the Department of Science and Innovation and the National Research Foundation (Grant No 98341) and the South African Medical Research Council. S.I.R. is supported by an H3 Africa grant (U01A136677) and the South African Medical Research Council. The funders had no role in the conceptualization or execution of any studies or drafting of the manuscript.

## AUTHOR CONTRIBUTIONS

Conceptualization and Methodology: M.J.V. and I.S.G.; Investigation: M.J.V., N.R., P.K., N.P. M., T.H., S.I.R., A.A.A., K.R.G., P.T.W., S.S., B.D., J.A., C.A.; Writing – Original Draft: M.J.V and I.S.G.; Writing – Review and Editing: All authors; Funding Acquisition: M.J.V. and I.S.G.; Resources: I.S.G.; Supervision: M.J.V., P.L.M. and I.S.G.

---

**Supplemental Figure 1:**
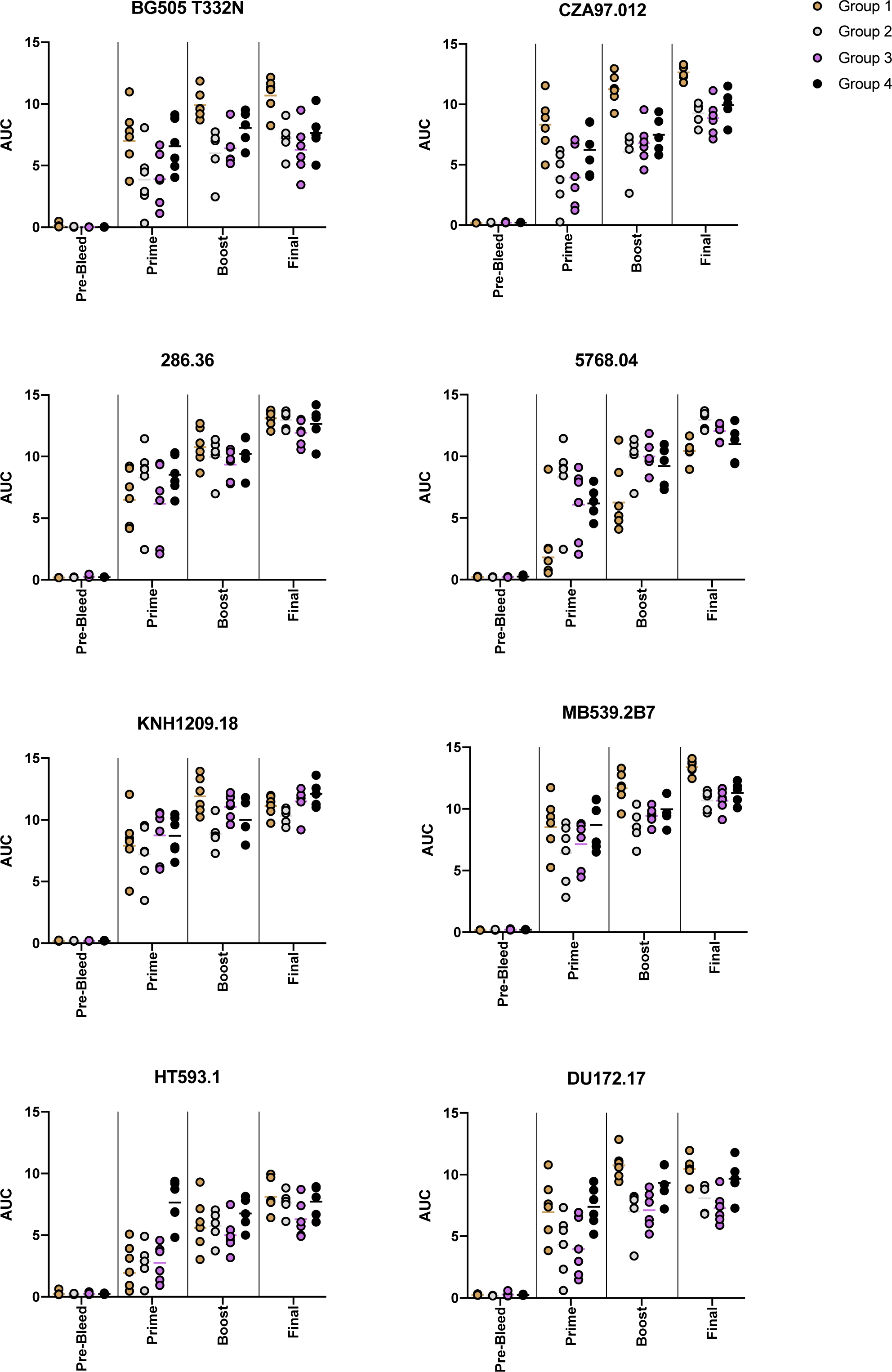
Serum ELISA binding. Sera from all timepoints were tested for binding against all strains of Env antigen in the vaccine regimen in an ELISA format. Final bleeds from all groups showed high binding to all Envs tested, whereas pre-bleed samples showed no binding. AUC, area under the curve.

**Supplemental Figure 2:**
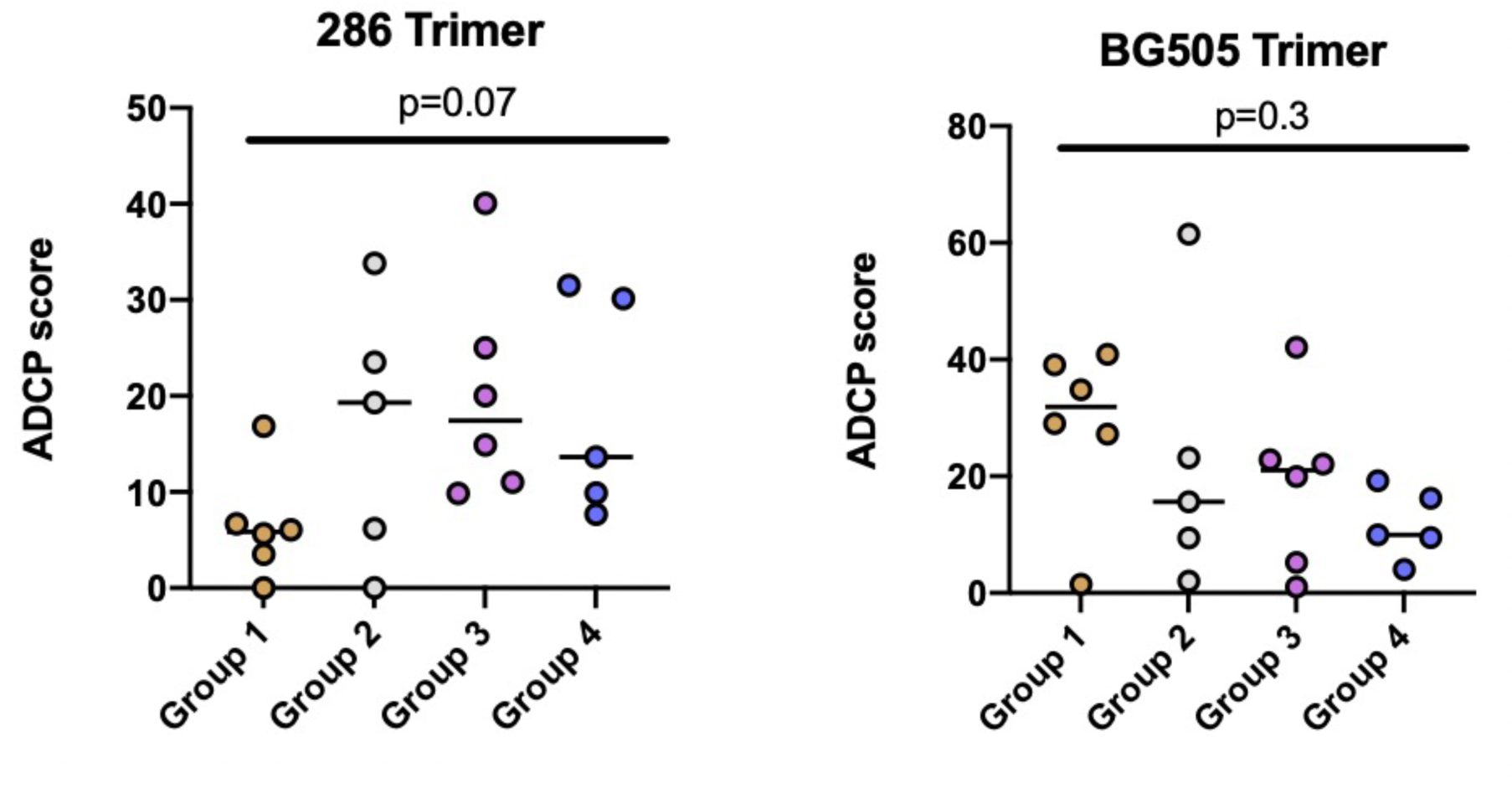
Antibody-Dependent Cellular Phagocytosis (ADCP) assay. Sera from all groups showed measurable ADCP responses to two strains of Env antigen. Fluorescent beads with immobilized streptavidin were coated with biotinylated Env trimer and subsequently mixed with sera from each guinea pig. THP-1 monocytes were then added and ADCP was quantified as the internalization or engulfment of the antigen coated beads by THP-1 cells measured by flow cytometry. This internalization is indicated as an ADCP score.

**Supplemental Figure 3:**
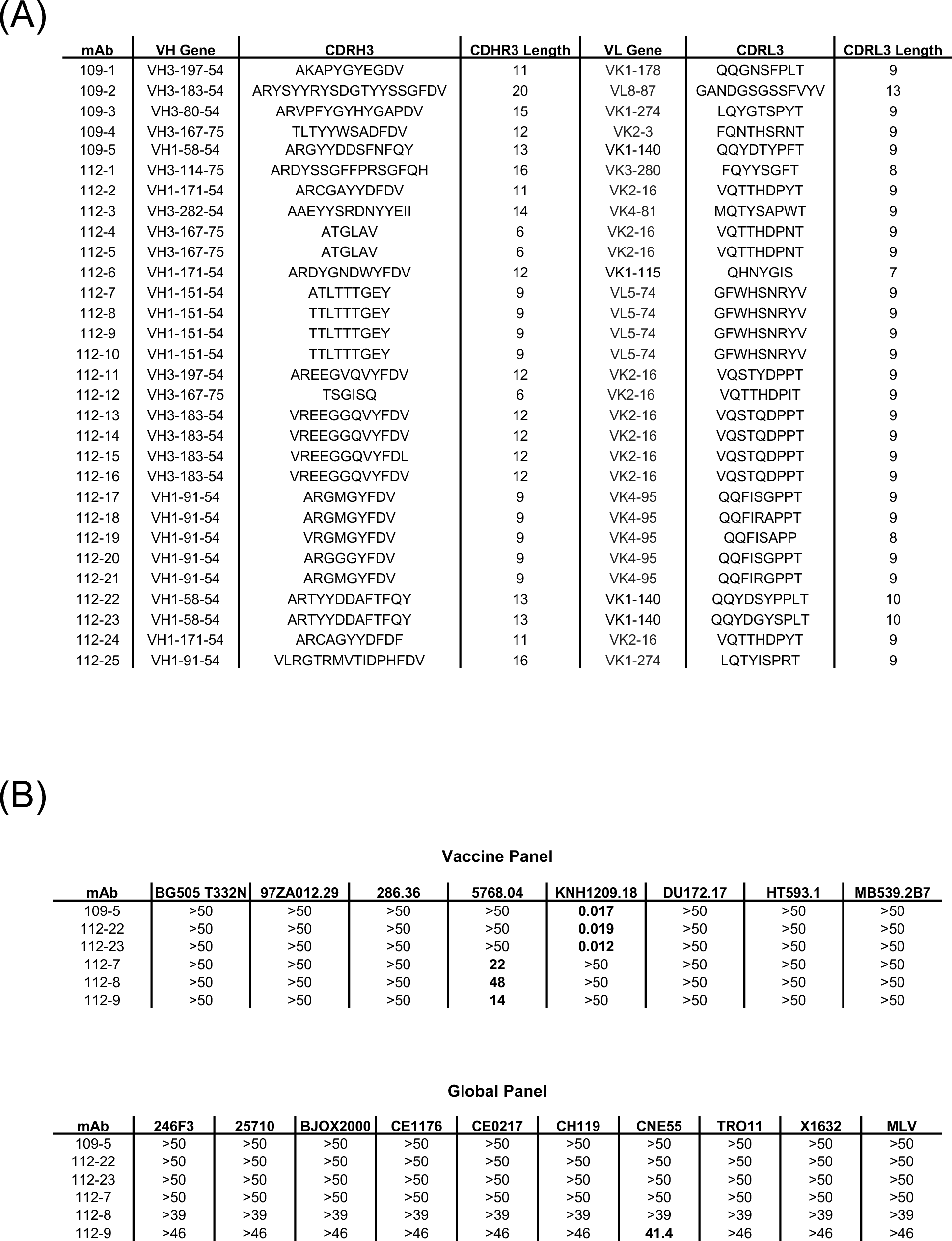
Guinea pig antibody gene usage and neutralization. 3A: Antibody sequences were aligned to published guinea pig V genes. CDR3 sequences are shown along with CDR3 lengths. 3B: Neutralizing antibodies were tested against the vaccine panel and 9-strain global panel. IC_50_ values are displayed in μg/mL. Bolded numbers indicate neutralization was detected.

**Supplemental Figure 4:**
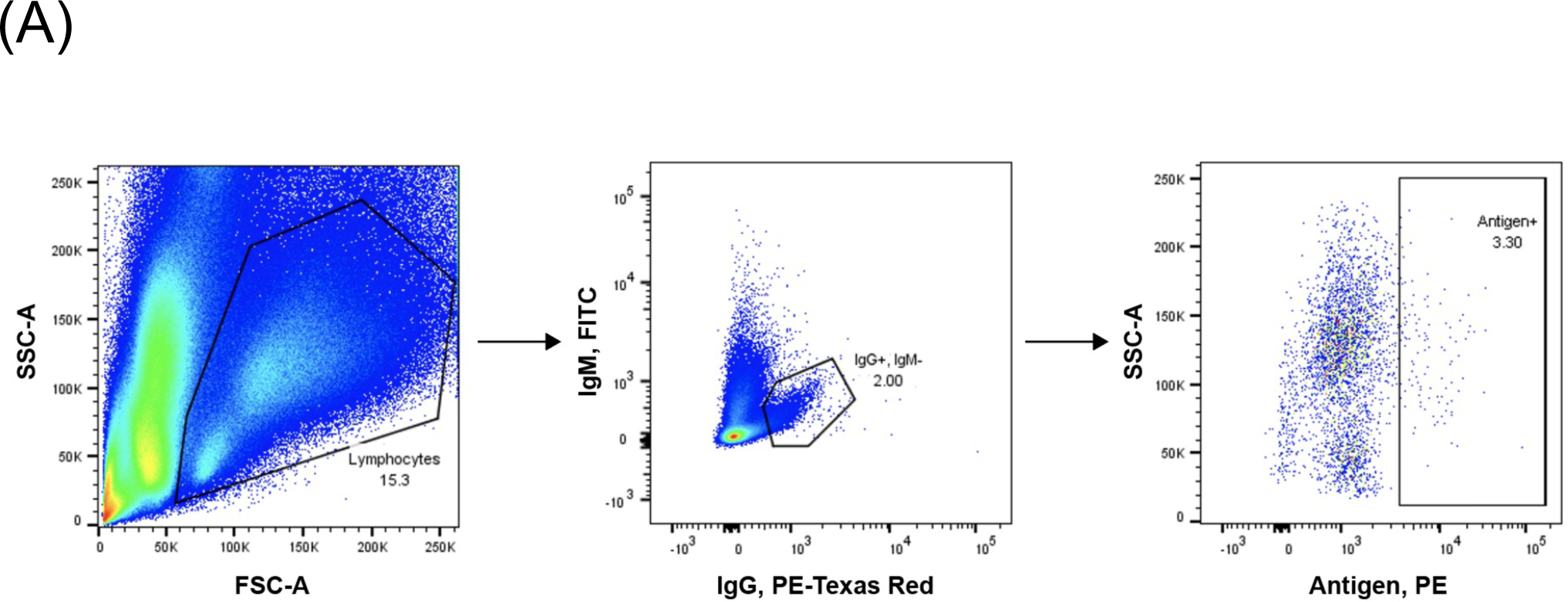
Flow data for GP109. Lymphocytes were gated out based on size, followed by live cells (data not shown). Cells with high IgG expression and negative for IgM expression were then gated followed by antigen positive B cells.

**Supplemental Figure 5:**
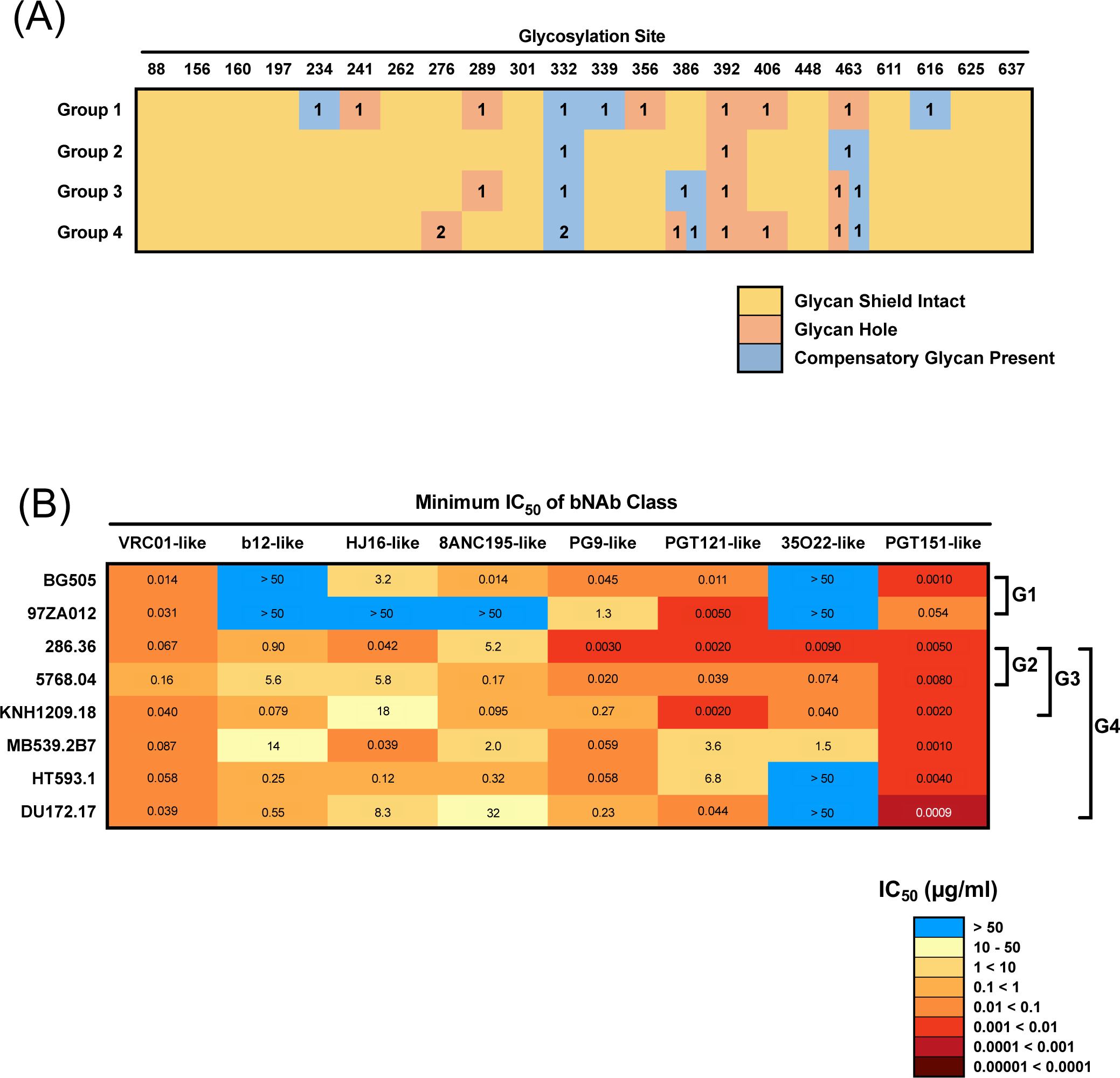
Vaccine groups 2-4 were selected to have intact glycan shields and high sensitivity to all major bNAb classes. 5A: Glycan occupancy at conserved glycosylation positions for each group is shown. Tan boxes indicate that all strains in a given group have a PNGS at the specified position. Pink boxes indicate a glycan hole, with the number denoting how many strains in the group are missing the specified glycan. Blue boxes denote a glycan hole that is compensated for by a neighboring glycan, with the number denoting how many strains in the group have the compensatory glycan. 5B: Vaccine strains are shown with the minimum IC_50_ values for each major bNAb class. Antibodies were grouped into bNAb classes based on epitopes targeted and other similarities such as angles of approach.

**Supplementary Figure 6:**
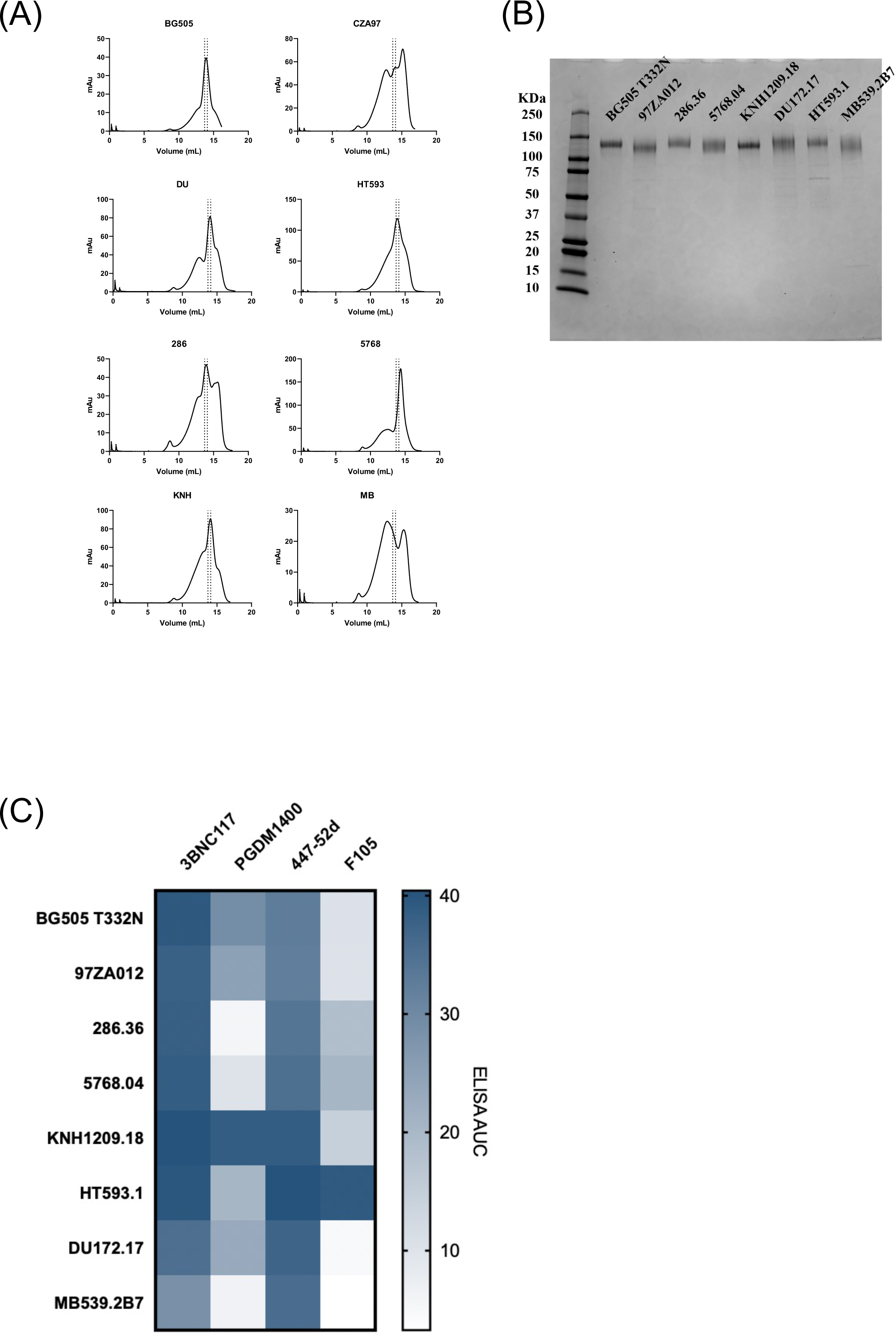
Immunogen production and QC. 6A: Env SOSIP.DS.664 immunogens displayed expected FPLC SEC profiles. Dotted lines indicate fractions collected indicative of trimers. 6B: Immunogens displayed expected size on SDS-PAGE (reducing conditions). 6C: Antigenicity of antigens was as expected measured by ELISA using a panel of standard antibodies including a CD4 binding site antibody (3BNC117), a quaternary-specific antibody (PGDM1400), as well as antibodies that recognize the open form of the trimer (F105 and 447-52d).

## MATERIALS AND METHODS

### HIV-1 Env strain selection algorithm

Prior work has established key factors related to the characteristics of Env strains used for immunization. These include the presence of an intact glycan shield at conserved positions, the availability of bNAb epitopes, the number of strains, as well as the level of amino acid sequence diversity between the strains, included in a multivalent vaccine. We developed a multi-objective optimization algorithm that selects for groups of Env strains that are optimized for the features listed above (Figure 1A and Figure S5). *Glycan Shield Coverage:* A set of ∼5,000 representative HIV-1 strains was selected from the Los Alamos National Laboratory HIV database and their Env amino acid sequences were aligned to the reference HXB2 strain. From the Env alignment, all residue positions that corresponded to an N-linked glycosylation sequon were extracted from each strain. Residue positions for which at least 50% of strains have an N-linked glycosylation sequon were defined as conserved glycan positions. For strains that were missing conserved glycans, structural analysis of the Env trimer structure was performed to identify potential compensatory glycans that are within 10Å of a missing conserved glycan. For each strain, the list of conserved and compensatory glycans was then used for further analysis. The completeness of the glycan shield is shown for each vaccine group (Figure S5A). *Availability of bNAb Epitopes:* The algorithm was designed to select Envs from HIV-1 strains that are sensitive to every major bNAb class, and thus present numerous epitopes that are capable of eliciting antibodies that may evolve neutralization breadth. Published datasets of bNAb-virus neutralization were used to divide bNAbs into discrete sets of epitope specificities (e.g. VRC01-like, PG9-like, etc.). For each strain and bNAb specificity group, the minimum (best), median, and maximum (worst) neutralization IC_50_ values among all bNAbs in that group were computed. Strains with minimum neutralization values of greater than 1 µg/ml for any bNAb specificity group, and strains for which two or more bNAb groups include an antibody that cannot neutralize the given strain (IC_50_ value of >50 µg/ml) were filtered out. The remaining strains were used for further optimization. Neutralization sensitivities of each strain composing our vaccine groups for every major bNAb class are shown (Figure S5B). *Env Sequence Diversity:* The Env sequence diversity within each combination of strains was computed. This was done both for the entire Env sequence (to account for overall clade diversity), as well as specifically for the protein surface residues (to account for antibody epitope diversity). The number of Env strains used in a multivalent vaccine may have opposing effects: on one hand, adding more strains may allow for closer mimicking of virus swarms during HIV-1 infection; on the other hand, the inclusion of more strains may increase the likelihood of generating off-target antibody responses. To assess this effect, we allowed the algorithm to pick set sizes of 2, 3, or 6 Env strains (Figure 1A).

### Immunogen expression and purification

We designed immunogens using the SOSIP platform to create soluble Env proteins stabilized in the prefusion conformation. In addition to the SOS (A501C and T605C) and IP (I559P) mutations that form the base of the SOSIP platform, we truncated Envs at position 664 and added the DS (I201C and A433C) mutation set to prevent CD4 triggering[27, 28]. The furin cleavage site between gp120 and gp41 was replaced with a serine-glycine linker (length 15) to create single-chain constructs [29]. The immunogens were produced in Expi293F cells by transient transfection in FreeStyle F17 expression media (Thermo Fisher) supplemented to a final concentration of 0.1% Pluronic Acid F-68 and 20% 4mM L-glutamine using Expifectamine transfection reagent (Thermo Fisher Scientific), cultured for 4-7 days at 8% CO_2_ saturation and 37°C with shaking. After transfection, cultures were centrifuged at 4000 g for 20 minutes.

Supernatant was filtered with Nalgene Rapid Flow Disposable Filter Units with PES membrane (0.45μm), and then run slowly over a column with agarose-bound Galanthus nivalis lectin (Vector Laboratories cat no. AL-1243-5) at 4°C. The column was washed with PBS, and protein was eluted with 15 mL of 1M methyl-α-D-mannopyranoside. The protein elution was buffer exchanged 3 times into PBS and concentrated using 100kDa Amicon Ultra centrifugal filter units. Concentrated protein was run on a Superose 6 Increase 10/300 GL on the AKTA FPLC system. Peaks corresponding to trimeric species were identified based on elution volume and antigenicity to quaternary structure-preferring antibody (PGDM1400) (Figure S6).

### Immunizations

Immunizations were conducted by, and in accordance to the guidelines of, Cocalico Biologicals, Inc., Reamstown, PA, USA. Immunizations were done at four week intervals, for a total of three injections per guinea pigs. A pre-bleed sample was taken prior to the first immunization. Guinea pigs were healthy, outbred, research-naïve, between 350-500 grams, about 1-2 months of age, half male and half female for each vaccine group. Immediately before each administration, 100 μL of potein mix (100 μg total Env) in PBS was mixed with 325 μL of CpG at 0.154 μg/μL (IDT), then with 75 μL of Emulsigen-D (MVP Adjuvants), giving a total volume of 0.5 mL. The final mixture was vortexed 5 times with 10 second intervals, and then was administered bilaterally in the quadricep muscles with half (0.25 mL) in the left quadricep and half (0.25 mL) in the right quadricep. Bleeds were obtained from the vena cava of anesthetized animals four weeks after each immunization. Spleens were collected for splenocyte isolation four weeks after the final immunization.

### Biotinylation of antigens

Protein antigens used for LIBRA-seq and serum ELISA contained a C-terminal Avi-tag and were site specifically biotinylated using BirA biotin-protein ligase raction kit (Avidity) according to manufacturer’s instructions.

### Oligonucleotide barcodes

We used oligos that possess a 15 bp antigen barcode, a sequence capable of annealing to the template switch oligo that is part of the 10X bead-delivered oligos and contain truncated TruSeq small RNA read 1 sequences in the following structure: 5’-CCTTGGCACCCGAGAATTCCANNNNNNNNNNNNNNNCCCATATAAGA*A*A-3’, where Ns represent the antigen barcode. Oligos were ordered from Sigma-Aldrich and IDT with a 5’ amino modification and HPLC purified. The following antigen barcodes were used: GCAGCGTATAAGTCA (CA/09), CAGTAAGTTCGGGAC (RSC3), TGACCTTCCTCTCCT (HT593.1), GACCTCATTGTGAAT (KNH1209.18), TAACTCAGGGCCTAT (5768.04), CAGATGATCCACCAT (286.36), ATCGTCGAGAGCTAG (BG505 T332N).

### Conjugation of oligonucleotide barcodes to antigens

For each antigen, a unique DNA barcode was directly conjugated to the antigen using a SoluLINK Protein-Oligonucleotide Conjugation kit (TriLink, S-9011) according to the manufacturer’s protocol. Briefly, the oligo and protein were desalted, and then the amino-oligo was modified with the 4FB crosslinker, and the biotinylated antigen protein was modified with S-HyNic. Then, the 4FB-oligo and the HyNic-antigen were mixed together. This causes a stable bond to form between the protein and the oligonucleotide. The concentration of the antigen-oligo conjugates was determined by a BCA assay, and the HyNic molar substitution ratio of the antigen-oligo conjuagtes was analyzed using the NanoDrop according to the Solulink protocol guidelines. AKTA FPLC was used to remove excess oligonucleotide from the protein-oligo conjugates, which were also checked using SDS-PAGE with a silver stain.

### Enrichment of antigen-specific guinea pig B cells

Splenocytes from each guinea pig were sorted and sequenced on separate days. Frozen splenocytes were thawed and resuspended in 10ml of pre-warmed RPMI 1640 medium supplemented with 10% heat-inactivated Fetal bovine serum (FBS). The cells were then washed with 10 ml of warm DPBS supplemented with 0.1% Bovine serum albumin (BSA). Cells were resuspended in DPBS-BSA and stained with cell markers including viability dye (Ghost Red 780), anti-guinea pig IgM-FITC, and anti-guinea pig IgG-Alexa Fluor 594 in the dark for 30 minutes at room temperature. Cells were then washed three times with DPBS-BSA at 300g for 5 minutes. Antigen-oligo conjugates were then added and incubated with the cells for 30 minutes in the dark. Cells were then washed three times with DPBS-BSA at 300g for 5 minutes. Streptavidin-PE was added to label the cells with bound antigen and incubated for 15 minutes in the dark at room temperature. Cells were then washed three times with DPBS-BSA, resuspended in DPBS-BSA, and then sorted by FACS.

### 10X Genomics single cell processing and next generation sequencing

Single-cell suspensions were loaded onto the Chromium Controller microfluidics device (10X Genomics) and processed using previously described methods for LIBRA-seq, with the exception that a custom primer set outlined in (Figure 3C) was used for target enrichment steps. Target enrichment step 1 consisted of a mix of Outer RP sequences 1-4 (final concentrations of 0.5μM each) and FP1 (final concentration 1μM). Target enrichment step 2 consisted of a mix of inner RP sequences 1-3 (final concentration of 0.5 μM each) and FP2 (final concentration of 1μM). A target capture of 10,000 B cells was used per 1/8 10X cassette for B cells. Slight modifications were made to intercept, amplify, and purify the antigen barcode libraries as previously described [26].

### Sequence processing and bioinformatics analysis

We followed our established pipeline, which takes paired-end FASTQ files of oligonucleotide libraires as input, to process and annotate reads for cell barcodes, unique molecular identifiers (UMIs) and antigen barcodes, resulting in a cell barcode-antigen barcode UMI count matrix [26]. B cell receptor contigs were processed using CellRanger 3.1.0 (10X Genomics) and GRCh38 Human V(D)J 7.0.0 as reference, while the antigen barcode libraries were also processed using CellRanger (10X Genomics). The cell barcodes that overlapped between the two libraries formed the basis of the subsequent analysis. Cell barcodes that only had non-functional heavy chain sequences as well as cells with multiple functional heavy chain sequences and/or multiple functional light chain sequences, were eliminated, reasoning that these may be multiplets. We also aligned the B cell receptor contigs (filtered_contigs.fasta file output by CellRanger, 10X Genomics) to IMGT references genes using IMGT/HighV-QUEST. The output of HighV-Quest was parsed using Change-O and combined with an antigen barcode UMI count matrix. Finally, we determined the LIBRA-seq score for each antigen in the library for every cell as previously described [26]. Raw sequencing results were aligned to the human VDJ reference genome using Cell Ranger to assemble heavy and light chains due to the absence of a suitable guinea pig VDJ reference. For guinea pig gene assignments of the antibodies shown in (Figure 4C) and (Figure S3), the immunoglobulin amino acid sequences for guinea pigs were obtained from [30] and aligned to amino acid sequences of antibodies discovered in our work described here using Clustal Omega.

### Public antibody analysis

Public clones were identified on the basis of a minimum of 70% amino acid sequence identity in both CDRH3 and CDRL3 regions and matching heavy and light variable (V) and joining (J) gene usage.

### Humanized guinea pig antibody expression and purification

The variable regions of guinea pig antibody sequences obtained from LIBRA-seq were appended into antibody expression vectors that contained human constant regions for heavy and light chains to create humanized guinea pig antibody sequences. Humanized guinea pig antibodies were produced in Expi293F cells by transient transfection in FreeStyle F17 expression media (Thermo Fisher) supplemented to a final concentration of 0.1% Pluronic Acid F-68 and 20% 4mM L-glutamine using Expifectamine transfection reagent (Thermo Fisher Scientific), cultured for 4-7 days at 8% CO_2_ saturation and 37°C with shaking. After transfection, cultures were centrifuged at 4000 g for 20 minutes. Supernatant was filtered with Nalgene Rapid Flow Disposable Filter Units with PES membrane (0.45μm), and then run slowly over a column with agarose-bound Protein A at 4°C. The column was washed with PBS, and antibody was eluted with 5 mL of 100mM glycine at a pH of 2.8. The elution was buffer exchanged 3 times into PBS and concentrated using 50kDa Amicon Ultra centrifugal filter units.

### Enzyme linked immunosorbent assay (ELISA) for immunogens

ELISA for immunogens utilized a lectin-based capture format. Galanthus nivalis lectin was plated at 2 μg/mL overnight at 4°C. The next day, plates were washing three times with PBS supplemented with 0.05% Tween20 (PBS-T) and blocked with 1% BSA in PBS-T. Plates were incubated at room temperature for one hour and then washed three times with PBS-T. Immunogens were diluted to 2 μg/mL in 1% BSA in PBS-T and then added to the plates. Plates were incubated at room temperature for two hours and then washed three times with PBS-T. A dilution series of antibodies was made in 1% BSA in PBS-T and then added to the plates. Plates were incubated at room temperature for one hour and then washed three times with PBS-T. The secondary antibody, goat anti-human IgG conjugated to peroxidase, was added at 1:10,000 dilution in 1% BSA in PBS-T to the plates, which were incubated for one hour at room temperature. Plates were washed three times and then developed by adding TMB substrate to each well. The plates were incubated at room temperature for five minutes, and then 1N sulfuric acid was added to stop the reaction. Plates were read at 450 nm. ELISAs were repeated two more times.

### ELISA for guinea pig serum

ELISA for serum was similar to as described above but with some modifications. Antigens with C-terminal Avi tags were site specifically biotinylated as described in above biotinylation section. Streptavidin was plated at 2 μg/mL overnight at 4°C. The next day, plates were washing three times with PBS supplemented with 0.05% Tween20 (PBS-T) and blocked with 1% BSA in PBS-T. Plates were incubated at room temperature for one hour and then washed three times with PBS-T. Biotinylated antigens were diluted to 2 μg/mL in 1% BSA in PBS-T and then added to the plates. Plates were incubated at room temperature for two hours and then washed three times with PBS-T. Serum was diluted starting at 1:50 (highest concentration) in 1% BSA in PBS-T, and then 5-fold down for a total of seven dilutions. Positive control antibodies for each antigen were included alongside. Plates were incubated at room temperature for one hour and then washed three times with PBS-T. The secondary antibody, goat anti-guinea pig IgG conjugated to peroxidase (highly cross-adsorbed), was added at 1:10,000 dilution in 1% BSA in PBS-T to the plates, which were incubated for one hour at room temperature. Plates were washed three times and then developed by adding TMB substrate to each well. The plates were incubated at room temperature for five minutes, and then 1N sulfuric acid was added to stop the reaction. Plates were read at 450 nm. ELISAs were repeated two more times.

### ELISA for humanized monoclonal guinea pig antibodies

ELISA for humanized monoclonal guinea pig antibodies was identical to ELISA for serum, with the exceptions that antibodies were diluted and screened (instead of serum), and the secondary was goat anti-human IgG conjugated to peroxidase (as used in ELISA for immunogens).

### Antibody-dependent cellular phagocytosis assay (ADCP)

The THP-1 phagocytosis assay was performed as previously described [31] using 1μM neutravidin beads (Molecular Probes Inc, Eugene, OR) coated with BG505 or 286 trimer. Sera were titrated and tested at a 1 in 100 dilution. Phagocytic scores were calculated as the geometric mean fluorescent intensity (MFI) of the beads multiplied by the percentage bead uptake as measured on a FACSAria II (BD biosciences, Franklin Lakes, New Jersey). THP-1 cells were obtained from the NIH AIDS Reagent Program and cultured at 37°C, 5% CO_2_ in RPMI containing 10% heat-inactivated fetal bovine serum (Gibco, Gaithersburg, MD) with 1% Penicillin Streptomycin (Gibco, Gaithersburg, MD) and not allowed to exceed 4 × 10^5^ cells/ml.

### TZM-bl neutralization assays

Serum and monoclonal antibody neutralization was assessed using the TZM-bl assay as described in [32]. This standardized assay measures antibody-mediated inhibition of infection of TZM-bl cells by molecularly cloned Env-pseudoviruses. Viruses that represent circulating strains (tier 2) were included. Murine leukemia virus (MLV) was included as an HIV-specificity control. Neutralization was measured as a reduction in luciferase gene expression after a single round of infection of TZM-bl cells in a 96 well plate. Results are presented as serum dilution required to inhibit 50% of virus infection (ID_50_), or as antibody concentration required to inhibit 50% of virus infection (IC_50_).

